# Mesoscale organization in the cell envelope of *Deinococcus radiodurans*

**DOI:** 10.1101/2022.01.29.478271

**Authors:** Domenica Farci, Patrycja Haniewicz, Dario Piano

## Abstract

S-layers are highly ordered coats of proteins localized on the cell surface of many bacterial species. In these structures, one or more proteins form elementary units that self-assemble into a crystalline monolayer tiling the entire cell surface. Here, the cell envelope of the radiation-resistant bacterium *Deinococcus radiodurans* was studied by high-resolution cryo-electron microscopy finding the crystalline regularity of the S-layer extended into the layers below. The cell envelope appears to be highly packed and resulting from a three-dimensional crystalline distribution of protein complexes organized in close continuity but allowing different degrees of voidness in the entire thickness. These insights grade S-layers to mesoscale hubs behaving as structural and functional architraves essential for the entire cell body.

## Introduction

Surface layers (S-layers) are ordered repetitions of proteinaceous units that tile the cell body of many prokaryotes (Sleytr *and* Glauert, 1975; Bahl *et al.*, 1997; Messner *et al.*, 1997). The periodical unit of S-layers consists of one or more proteins that, in some species, are further organized in multi-protein complexes (Fagan and Fairweather 2014; Farci *et al.*, 2018a; Farci *et al.*, 2021). These coating structures are common in Bacteria and Archaea and have essential roles in providing cell rigidity, shape, adhesion, and resistance to extreme conditions (Beveridge *et al.*, 1997; Pavkov-Keller *et al.*, 2011; Farci *et al.*, 2018b; Kumar *et al.*, 2021). Despite their relevance in the cell envelope, detailed molecular and *in situ* studies on S-layers are still limited leading to a distinct degree of uncertainty about their roles and functions. However, the self-assembly properties of S-layer proteins (Pum *et al.*, 2013), and the levels of energy consumption implied their high expression and displacement, self-stand for a deep functionalization and specialization of these structures. Being the forefront of the cell to the environment, S-layers represent one of the cell’s compartments more exposed to selective pressure and, in extremophiles, cope with harsh conditions that are typically deviating from the canonical ones sustaining life (Farci *et al.*, 2016). This fact ranks S-layers as one of the cell regions most subjected to specialization in bacteria.

One of the model organisms for S-layer studies is the radiation-resistant bacterium *Deinococcus radiodurans.* Its S-layer has been investigated by several techniques and its surface structure and porous organization were previously described at a low resolution (Baumeister *et al.*, 1982; Baumeister *et al.*, 1986; Müller *et al.*, 1996; Lister and Pinhero, 2001). In the last decade, the research done on this S-layer provided important details on its organization, composition, and properties, eventually resulting in an overall vision with features somehow overlooked since then. In particular, this S-layer was found to be composed of several proteins further organized into different protein complexes (Farci *et al.*, 2014; Farci *et al.*, 2015). Taken together, these studies provided multiple and indirect evidence suggesting the absence of a distinct separation between the S-layer and the underlying membranes (Rothfuss *et al.*, 2006; Farci *et al.*, 2014; Yu *et al.*, 2017; Farci *et al.*, 2020; Farci *et al.*, 2021). To shed a light on the S-layer extension and interaction with the rest of the cell envelope, here we explore the *in-situ* 3D organization of this S-layer. Cell envelope patches were characterized by the mean of 3D cryo-electron crystallography (cryo-EC) cross-referenced by cryo-electron tomography (cryo-ET) and subsequent subtomogram average. Results describe an unexpectedly sophisticated crystalline regularity, with a 3D organization consisting of an ordered juxtaposition of three multi-protein complexes recently observed in their 2D organization (Farci *et al.*, 2021) and here shown to extend into the entire cell envelope thickness. The periodical organization of these complexes results in a highly packed cell envelope structure that, despite the close continuity of its components, still allows for different degrees of voidness in the cell envelope thickness. While one of these complexes appears to be exclusive of the cell envelope surface, involving the S-layer and the outer membrane, the other two complexes, a Type IV-like Piliation system (T4P-like) and the S-layer Deinoxanthin Binding Complex (SDBC), that were previously described as channels (Gold *et al.*, 2015; Farci *et al.*, 2021), here are shown to span the layers below representing the main in/out system of sieving in the cell. Such an organization force the cell envelope trafficking through the two main available ways, the T4P-like and the SDBC, opening for a centralized and discrete functional displacement along and across the cell envelope. The presented findings candidate S-layers as fundamental structures that “drive” the cell envelope organization from the level of cell body down to molecular level according to a mesoscale model where active and central functions are implied.

## Results

### An extended crystalline organization characterizes the cell envelope of *Deinococcus radiodurans*

Given the general properties of radiation resistance and the abilities of its S-layer to shield the UV radiation (Daly, 2009; Farci *et al.*, 2016), direct radiation damage in *D. radiodurans* is likely to be damped by the robustness of its structures. This property makes *D. radiodurans* an optimal candidate for studies of the cell envelope by electron microscopy. Taking advantage of the S-layer crystallinity, the diffraction properties of the cell envelope were used as a discriminant for selecting the best patches to be subjected to cryo-EC, cryo-ET, and subtomogram average. Accordingly, only integral cell envelope patches showing both membranes and sharp diffraction patterns with reflections beyond 2.8 Å were selected (Fig. 1). Diffraction analyses by independent processing and subsequent merging of 73 micrographs at different tilting angles of six different patches led to a 3D electron density map that allowed an indirect visualization *in situ* of the crystalline regions in the cell envelope (Fig. 2; EMDB-14097). The phase residuals post-processing indicated information up to 4 Å (Table 1) and, interestingly, the reflections carried information not only of the S-layer, well known for its ~10 Å thick crystalline packing (Baumeister *et al.* 1986) but also of the underlying space, providing striking evidence of a crystallinity extended for ~200 Å into the cell envelope (Fig. 2 and Sup. Movie 1).

**Figure 1:**
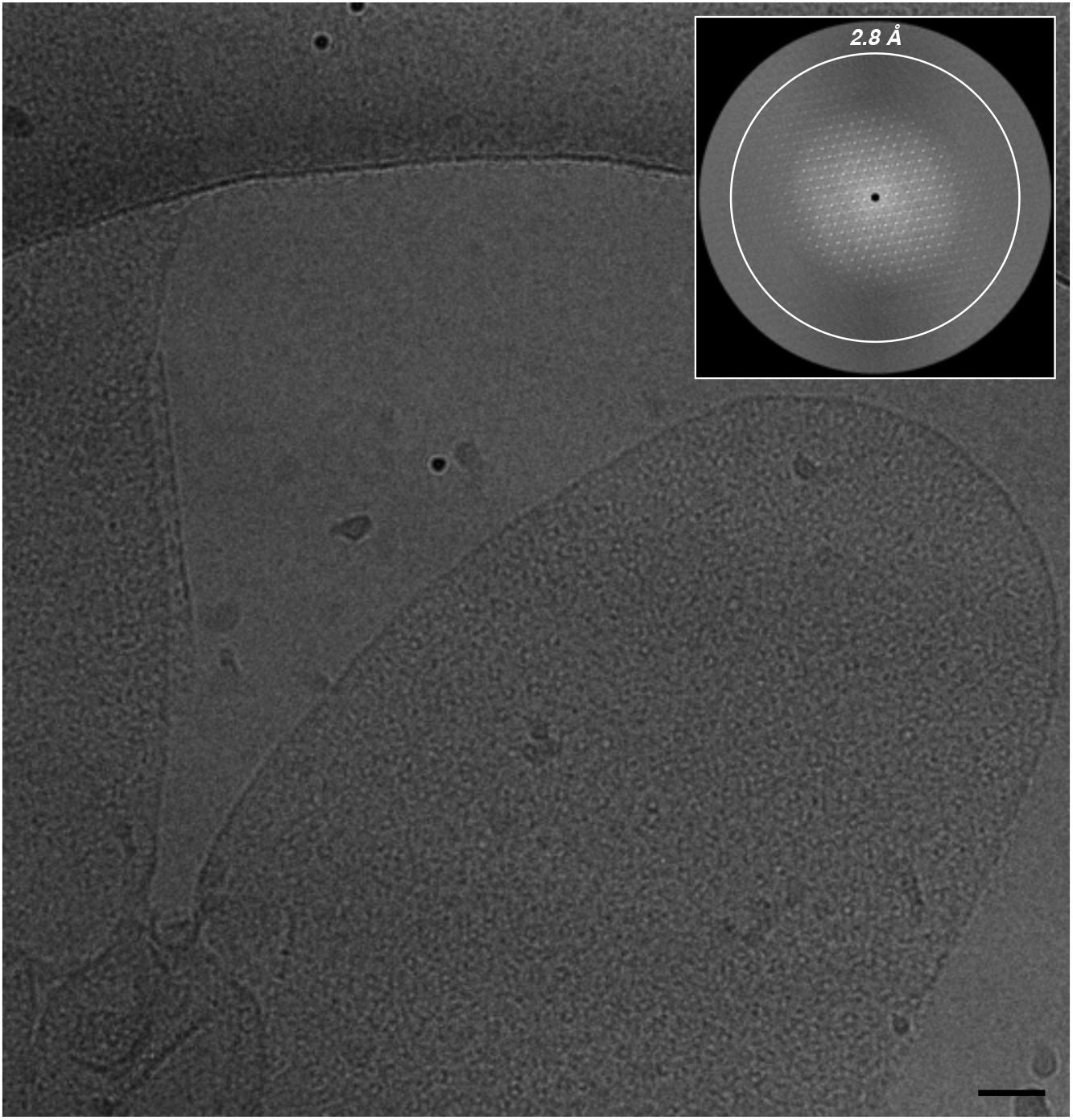
Continuous regularity of isolated cell envelope patches. Micrographs of cell envelope patches let glimpse a patterned regularity resulting from a crystalline organization. The crystalline nature of the sample can be visualized by electron diffraction with a pattern of diffraction spots beyond 2.8 Å resolution (inset). The scale bar indicates 500 Å.

**Table 1:**
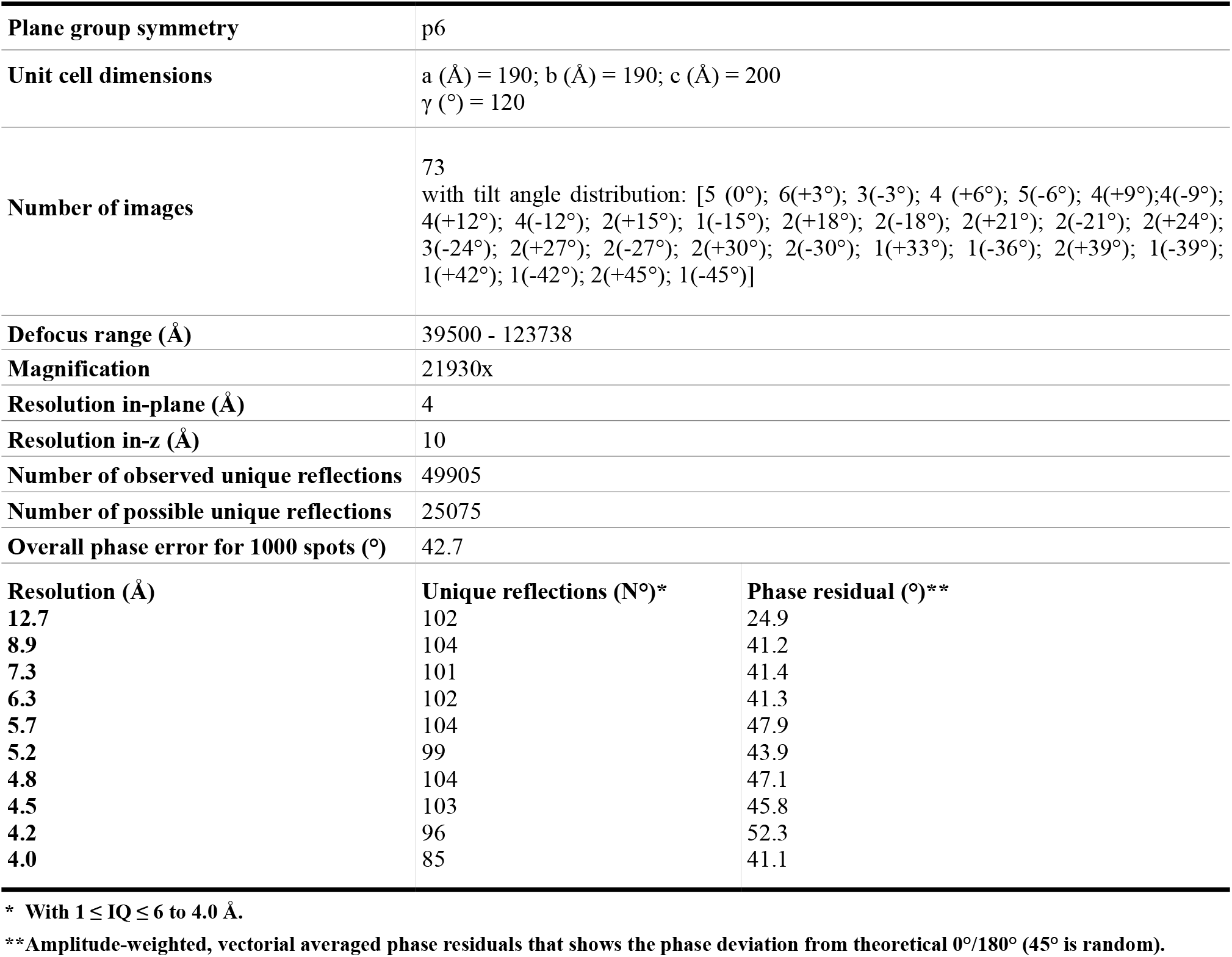
Electron crystallographic parameters and data.

|  |  |
| --- | --- |
| **Plane group symmetry** | p6 |
| **Unit cell dimensions** | a (Å) = 190; b (Å) = 190; c (Å) = 200<br>$\gamma$ (°) = 120 |
| **Number of images** | 73<br>with tilt angle distribution: [5 (0°); 6(+3°); 3(-3°); 4 (+6°); 5(-6°); 4(+9°); 4(-9°); 4(+12°); 4(-12°); 2(+15°); 1(-15°); 2(+18°); 2(-18°); 2(+21°); 2(-21°); 2(+24°); 3(-24°); 2(+27°); 2(-27°); 2(+30°); 2(-30°); 1(+33°); 1(-36°); 2(+39°); 1(-39°); 1(+42°); 1(-42°); 2(+45°); 1(-45°)] |
| **Defocus range (Å)** | 39500 - 123738 |
| **Magnification** | 21930x |
| **Resolution in-plane (Å)** | 4 |
| **Resolution in-z (Å)** | 10 |
| **Number of observed unique reflections** | 49905 |
| **Number of possible unique reflections** | 25075 |

|  |  |  |
| --- | --- | --- |
| **Overall phase error for 1000 spots (°)** | 42,7 |  |
| **Resolution (Å)** | **Unique reflections (N°)\*** | **Phase residual (°)\*\*** |
| 12.7 | 102 | 24.9 |
| 8.9 | 104 | 41.2 |
| 7.3 | 101 | 41.4 |
| 6.3 | 102 | 41.3 |
| 5.7 | 104 | 47.9 |
| 5.2 | 99 | 43.9 |
| 4.8 | 104 | 47.1 |
| 4.5 | 103 | 45.8 |
| 4.2 | 96 | 52.3 |
| 4.0 | 85 | 41.1 |

\* With $1 \leq IQ \leq 6$ to 4.0 Å.
\*\*Amplitude-weighted, vectorial averaged phase residuals that shows the phase deviation from theoretical 0°/180° (45° is random).

**Table 1** - Electron crystallographic parameters and data
|  |  |  |
| --- | --- | --- |
| **Plane group symmetry** | p6 |  |
| **Unit cell dimensions** | a (Å) = 190; b (Å) = 190; c (Å) = 200<br>$\gamma$ (°) = 120 | |
| **Number of images** | 73<br>with tilt angle distribution: [5 (0°); 6(+3°); 3(-3°); 4 (+6°); 5(-6°); 4(+9°);4(-9°); 4(+12°); 4(-12°); 2(+15°); 1(-15°); 2(+18°); 2(-18°); 2(+21°); 2(-21°); 2(+24°); 3(-24°); 2(+27°); 2(-27°); 2(+30°); 2(-30°); 1(+33°); 1(-36°); 2(+39°); 1(-39°); 1(+42°); 1(-42°); 2(+45°); 1(-45°)] |  |
| **Defocus range (Å)** | 39500 - 123738 |  |
| **Magnification** | 21930x |  |
| **Resolution in-plane (Å)** | 4 |  |
| **Resolution in-z (Å)** | 10 |  |
| **Number of observed unique reflections** | 49905 |  |
| **Number of possible unique reflections** | 25075 |  |
| **Overall phase error for 1000 spots (°)** | 42.7 |  |
| **Resolution (Å)** | **Unique reflections (N°)\*** | **Phase residual (°)\*\*** |
| 12.7 | 102 | 24.9 |
| 8.9 | 104 | 41.2 |
| 7.3 | 101 | 41.4 |
| 6.3 | 102 | 41.3 |
| 5.7 | 104 | 47.9 |
| 5.2 | 99 | 43.9 |
| 4.8 | 104 | 47.1 |
| 4.5 | 103 | 45.8 |
| 4.2 | 96 | 52.3 |
| 4.0 | 85 | 41.1 |

\* With $1 \leq IQ \leq 6$ to 4.0 Å.
\*\*Amplitude-weighted, vectorial averaged phase residuals that shows the phase deviation from theoretical 0°/180° (45° is random).

**Figure 2:**
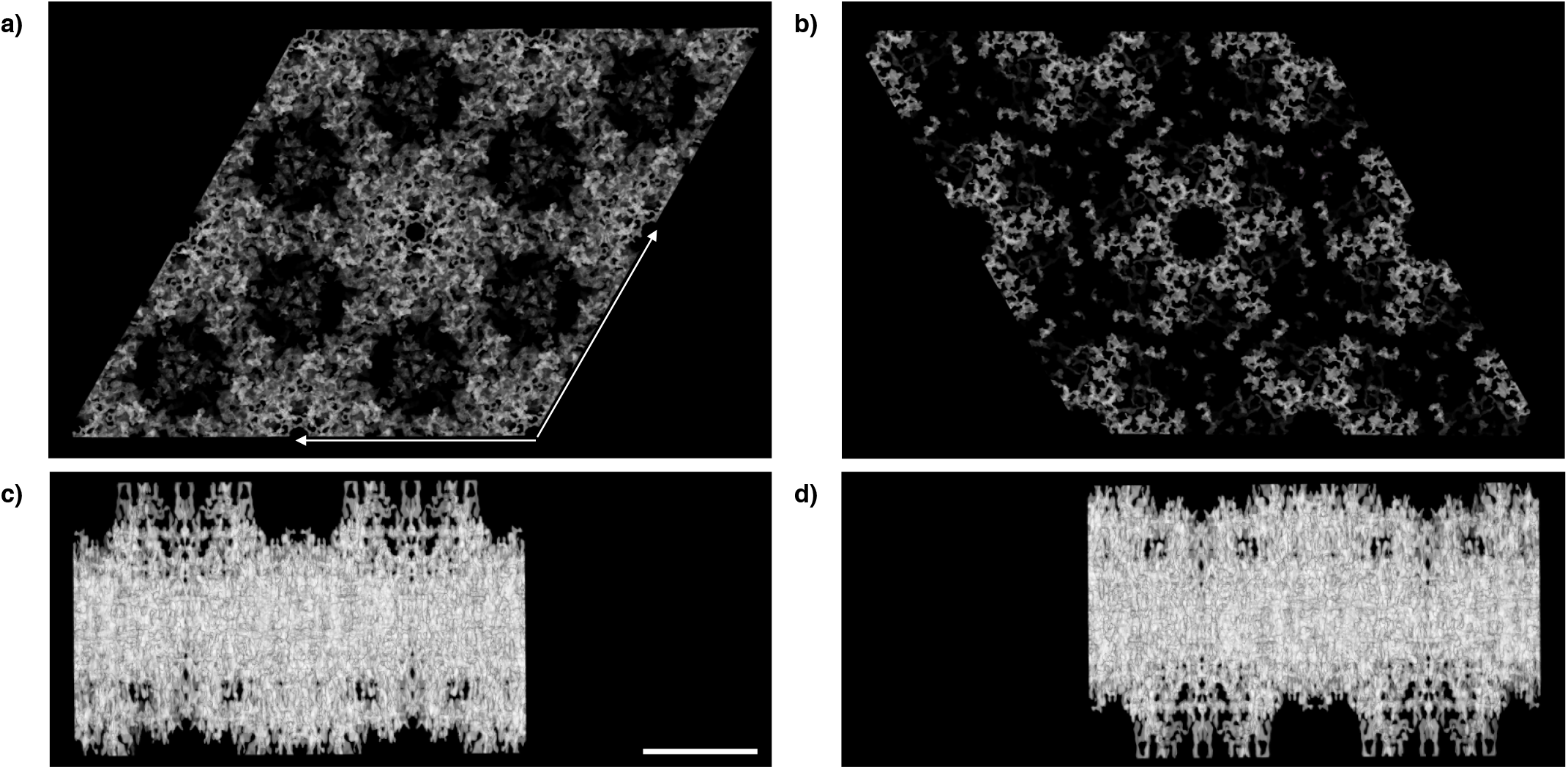
Projection structures of the crystalline cell envelope fraction. Map at 4 Å with imposed p6 symmetry (EMDB-14097) visualized on the top view (a), bottom view (b), and side views (c and d). The map clearly shows the main structural constituents organized in individual units. In the side views, the thickness reaches ~200 Å. The scale bar indicates 100 Å.

### Three different protein complexes account for the three-dimensional cell envelope crystalline packing

A consistent volume of the cell envelope appears to be occupied by three complexes. The larger complex shows an overall shape that resembles one of many secretion systems and fits well with the ~1.1 MDa complex, called Type IV Piliation-like system (T4P-like; Fig. 3a) previously identified. This complex was found to be abundant in this cell envelope and to have as a main component the protein DR_0774, a pilin Q commonly constituting the head region and interacting with the outer membrane in many secretion systems (Farci *et al.*, 2014). With a height of 175 Å (Fig. 3a), a main hexagonal head with a side of 85 Å and a diameter of 150 Å (Fig. 3b, d, and g), the T4P-like complex spans the S-layer, the outer membrane, and the periplasm reaching the inner membrane (Fig. 3a and f). On the top of the head, a multiple-spikes crown emerges and expands for 20 Å defining the pore region (Fig. 3a). According to the p6 symmetry of this S-layer, each T4P-like is surrounded by 6 copies of a smaller complex, the S-layer Deinoxanthin Binding Complex (SDBC), extensively described and characteristic of these cell envelopes (Farci *et al.*, 2016; Farci *et al.*, 2019; Farci *et al.*, 2020; Farci *et al.*, 2021). Recognizable by its triangular shape (with a side length of 90 Å and a thickness of ~30 Å), this ~0.9 MDa complex has the main body embedded in the outer membrane and is located inside of a “cell envelope case” (Fig. 3b, c, d, and f), as already observed in middle-high resolution studies on the surface of this cell envelope (Farci *et al.*, 2021). Differently from what was observed before in purified SDBC samples (Farci *et al.*, 2021), here from the main body depart three harms facing the external environment and the other three toward the periplasmic side (Fig. 3c and f). Finally, a third dihedral complex of unknown identity, characterized by a typical C2 symmetry with a side of 70 Å, a width of 45 Å (Fig. 3b, d, and g), and a height of 60 Å (Fig. 3a and f), was found to be localized in the S-layer and to span the underlying outer membrane (Fig. 3a). By exclusion, we hypothesize that this third complex could be the DR_2508 protein, which is known as HPI (Hexagonally Packed Intermediate) (Baumeister *et al.,1982;* Baumeister *et al.*, 1986; Rothfuss *et al.*, 2006) and is the only known protein of this S-layer for which a precise localization is still missing. These three individual units, the T4P-like, the SDBC, and the dihedral complex here represented in orange, pink, and yellow, respectively, were extracted from the map providing a better view of their features and relative localization (Fig. 3f, g, and h; Sup. Movie 1).

**Figure 3:**
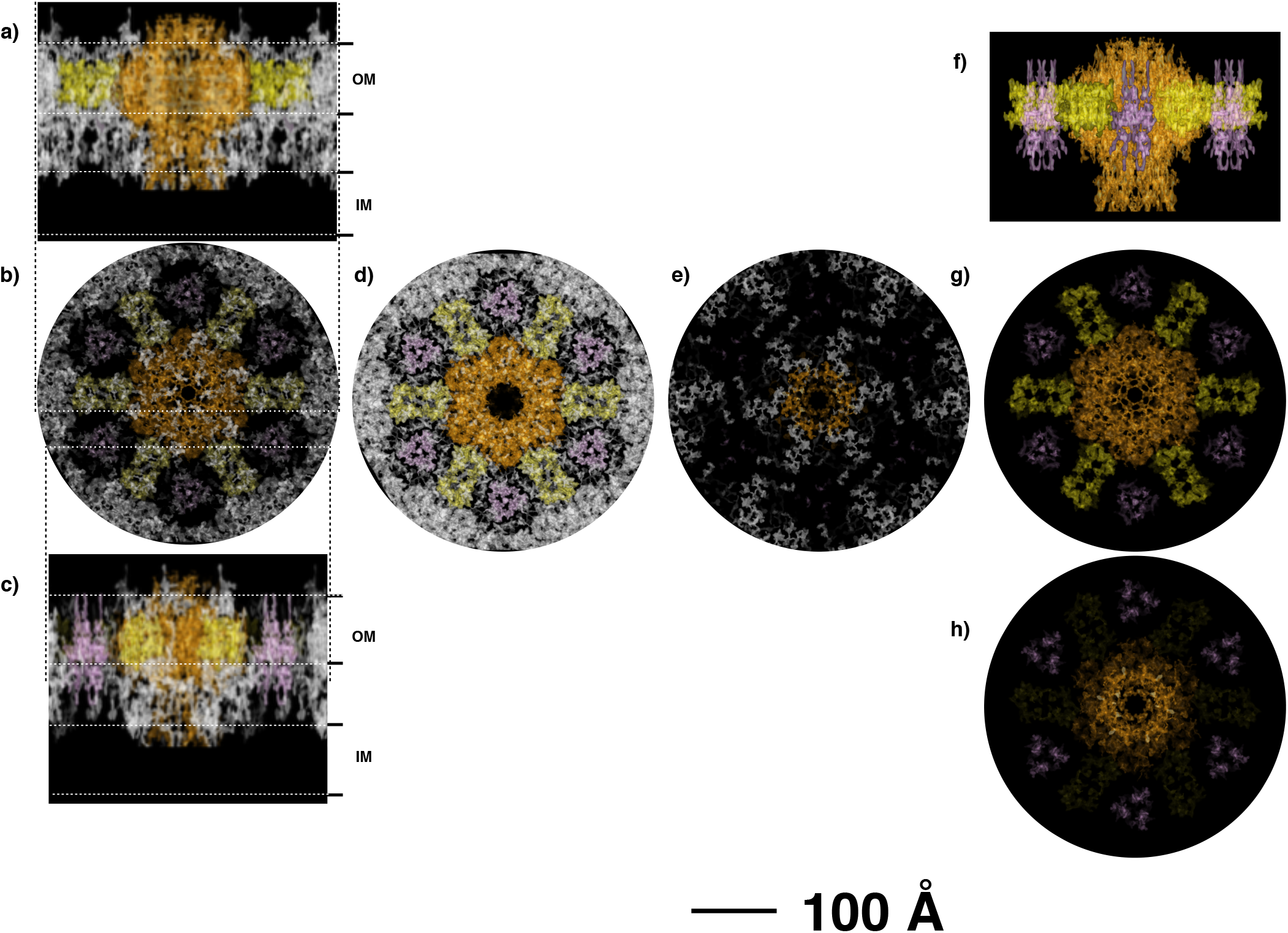
Projection structure with assigned densities. The specific occupancies for each complex are shown as side views at two sliced levels allowing to visualize the T4P-like in orange and the dihedral complex in yellow (a), and the SDBC in pink (c). For a better visualization slices at the top view level (b), at the outer membrane level (d), and at the bottom level (e) of the map are shown. Colored complexes represent the extracted densities from the original map. Side (f), top (g), and bottom (h) views of the extracted complexes and their relative position are also shown. The scale bar indicates 100 Å.

As a preliminary cross-checks of these observations, the mild solubilization of the cell envelopes sample and its subsequent fractionation by anionic exchange chromatography confirmed the dominance of these three protein complexes (Fig. 4a), as eventually shown by negative stain electron microscopy (Fig. 4b, c, and d).

**Figure 4:**
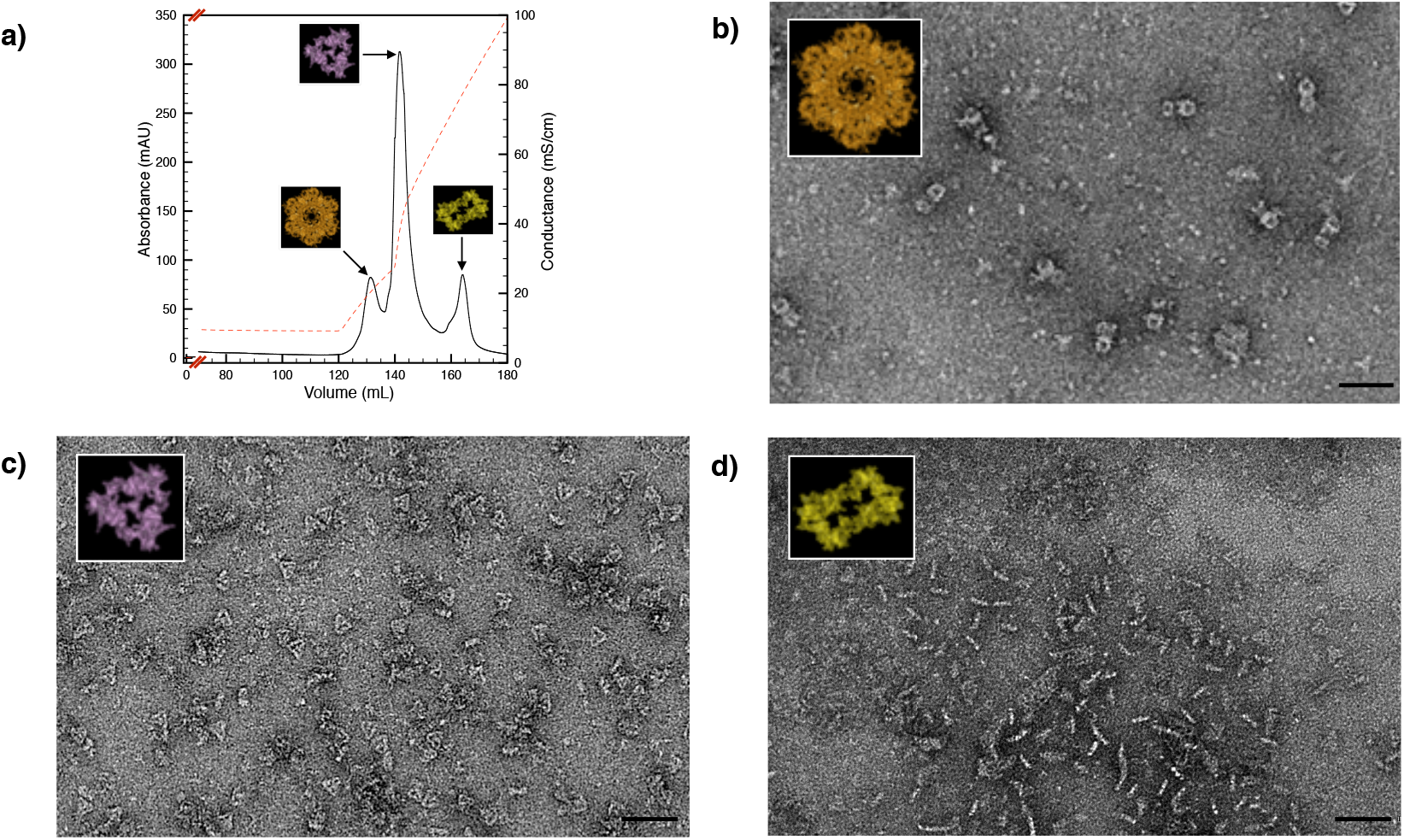
Chromatographic separation of the three cell envelope complexes. The cell envelope patches were mild solubilized and subsequently subjected to Anionic Exchange Chromatography (a) resolving in three main protein complexes: the T4P-like complex (b), the SDBC (c), and the dihedral complex (d). The scale bar indicates 500 Å.

### Cryo-electron tomography and subtomogram average on intact cell envelopes display an ordered repetition of channels confirming the cryo-EC analysis

Cryo-ET and subtomogram average on the same data sets allowed an *in-situ* direct analysis of intact cell envelope fragments (Fig. 5; EMDB-14095). At the tomographic level, raw movies’ slices (Fig. 5) and reconstructions (Sup. Movie 2) showed a repetition of proteinaceous units as top and side views. Overall, these analyses confirmed the organization described by cryo-EC, in particular showing a regular isoporous surface (interpore distance of 195 Å vs 190 Å measured by 3D cryo-EC) with each pore corresponding to a T4P-like complex spanning the whole layers of the cell envelope (Fig. 5, upper-middle and upper-right insets, respectively), hence granting for *in situ* direct visualizations of the 3D regularity. A subsequent p6 symmetrization centred in the pore region allowed to better resolve the T4P-like complex (8 Å best resolution; Fig. 6, Sup. Fig. 1, and Sup. Fig. 2; EMDB-14096) and compare it with the T4P-like structure extracted from the 3D cryo-EC map (Fig. 6). The subtomogram shows a clear dominance of the T4P-like occupancy (Sup. Fig. 2) with each complex having a side of 85 Å, a diameter of ~150 Å, and a total height of ~260 Å (Fig. 6). Included in this structure are also non-crystalline regions that are missing in the model obtained by cryo-EC. Noteworthy there are a 45 Å crown, which is more extended with respect to the one visible in the 3D diffraction data (~20 Å), and a 70 Å height conical extension in the bottom part of the complex involving the inner membrane and the neighbouring cytoplasmic region (Fig. 6, Sup. Fig. 2). Furthermore, slicing along the complex’s height allowed to compare the patterning of symmetry at different levels confirming the consistency of the two independent analyses (Fig. 6, left and right boxes).

**Figure 5:**
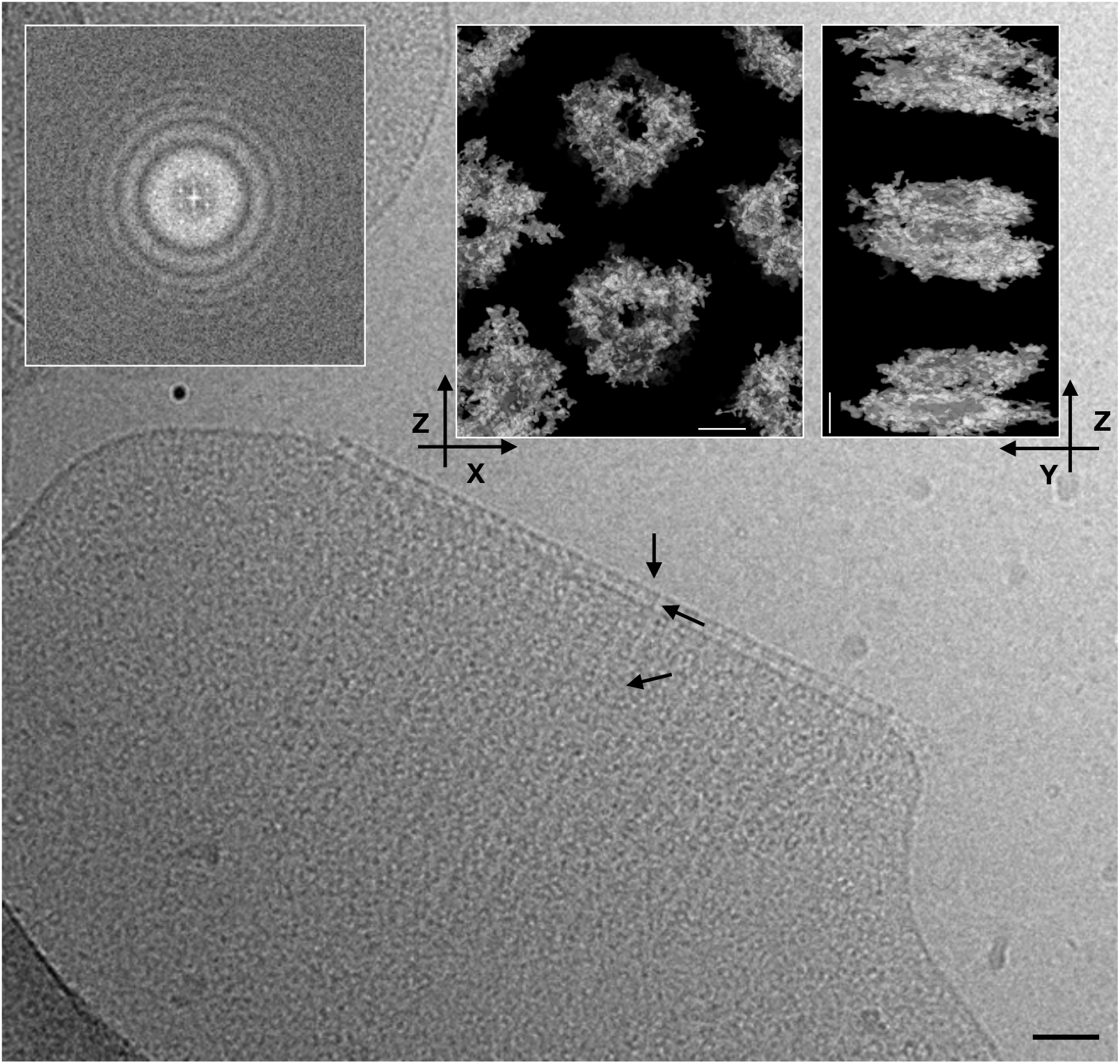
Tomography and related subtomogram average analysis. a) Slice of a raw tomographic movie and FFT diffraction (upper-left inset) on a cell envelope patch. A scale bar of 500 Å is shown. In the inset, it is shown a typical subtomogram average of cell envelope patch (EMDB-14095) from the top view (upper-centered inset) and side view (upper-right inset). No symmetry was imposed. The scale bar indicates 50 Å.

**Figure 6:**
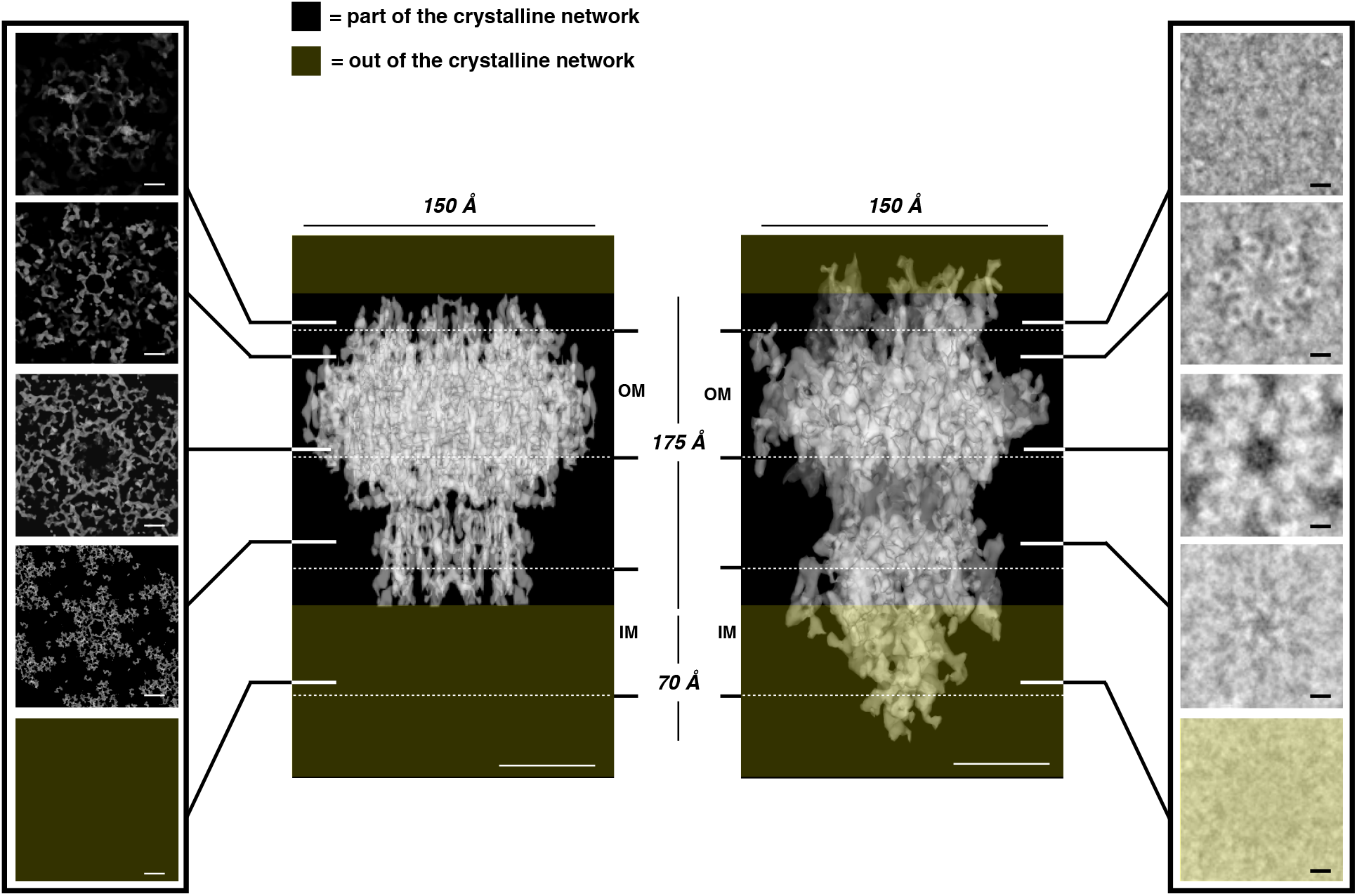
Features comparison between tomographic and crystallographic T4P-like. The main dimensions of the T4P-like complex obtained by 3D electron diffraction (center left; EMDB-14097) and by subtomogram average (center right; EMDB-14096) are compared. The left and right boxes show comparable images of the slice through at equivalent levels, from top to bottom, for the T4P-like complex obtained by cryo-EC (left box) and by cryo-ET subtomogram average (right box). Both maps were obtained by computing with imposed a p6 symmetry. Scale bars indicate 50 Å.

Finally, this analysis also allowed to measure the thickness of the entire cell envelope and its regions. The whole thickness was found to be of 299.5 ± 2.8 Å with a periplasm ~99.3 ± 12.2 Å thick delimited by a compact “roof” represented by the S-layer and the outer membrane tightly interacting to each other for ~98.9 ± 9.4 Å, and below by a “floor” consisting of an inner membrane ~69.8 ± 5.4 Å thick (Sup. Fig. 3).

## Discussion

We provide here the supramolecular description of the cell envelope in the radiation-resistant bacterium *D. radiodurans.* The natural capability of this organism in withstanding high doses of radiations along with the known crystalline properties of its S-layer were advantageous to perform a detailed characterization of the cell envelope. The intrinsic properties of radiation resistance most likely delayed the electron damage of the samples resulting in a well-preserved overall structure and allowing a middle-resolution characterization by 3D cryo-EC (Fig. 1 and 2) cross-referenced by cryo-ET (Fig. 5).

We demonstrate that the well-known S-layers’ order in this specific case is ascribable to three multi-protein complexes (Fig. 3). The most astonishing and remarkable result is that the S-layer regularity pervades the underlying membranes, imposing a defined order to the cell envelope (Fig. 3). The intricate structure resulting from a regular juxtaposition of protein complexes is expected to serve as an extended molecular sieving machine. This observation is unequivocally supported by its constituents, the T4P-like and the SDBC, already known for their gating properties (Gold *et al.* 2015; Farci *et al.* 2021). Accordingly, through this organization, the trafficking across the cell envelope has the potential to be finely regulated by the discrete distribution of its components. This complex simplicity, consisting of two types of gates isotropically distributed on the whole surface of the cell envelope, allows to equalized the movements in/out of the cell with respect to the surrounding environment. The observation of this regularity appeared first by the diffraction properties of the cell envelope patches and after was confirmed by the 3D structural analyses (Figs. 1 and 2).

Taken together the present findings suggest that the S-layer and the underlying membranes occur according to a clear mesoscale organization. In fact, as a consequence of this cell envelope topology, a continuous proteinaceous system can be envisaged from the range of individual cells down to molecular identities (Goodsell *et al.*, 2020). In such an organization, the crystalline unit cell represents a discrete structural unit, the repetition of which results in the association of 6 SDBC and 6 dihedral complexes for each T4P-like (Fig. 3). According to this spatial displacement and the self-assembling properties of S-layer proteins (Pum *et al.*, 2013), these three building blocks are expected to self-organize forming a large functional structure with several hierarchic levels, a characteristic feature of a mesoscale organization (Goodsell *et al.*, 2020). If on one hand, this structural continuity provides extended compactness and statical isotropy at different levels, on the other hand by this structure the bacterium has an isotropic interaction with the surrounding environment by reaching an equally distributed exchange potential. Considering the well-known role of secretion systems and porins, such as the T4P-like (Gold *et al.* 2015; Craig *et al.*, 2019) and the SDCB (Achouak *et al.*, 2001; Vergalli *et al.*, 2020; Farci *et al.* 2021), in exchanging across the cell envelope, this property is relevant not only for the dynamics associated with the trafficking across but also with the specific pool of functions that might specialize a given S-layer.

Notably, while the cell envelope compactness appears to be an intrinsic property due to its organization, our data show a significant free space with limited hindrances (Fig. 3a and f). This is particularly true for the periplasm where, trafficking processes and proteins not involved in the paracrystalline organization, need to have enough space for the functional activities related to cell maintenance and homeostasis (e.g., osmotic and energetic balance).

At this stage, it is not given to know how much this organization is common among different S-layers. Considering that similar symmetries and surface organizations are also associated with other S-layers, this extended cell envelope regularity could be a frequent feature for S-layers of isoporous type. These findings provide a new breath for studying S-layers and understanding their active roles in the physiology of bacterial cells.

## Materials and Methods

### Cell culturing

*Deinococcus radiodurans* cultures (strain R1; ATCC 13939) were cultivated in Tryptone Glucose Yeast extract broth (TGY) at 30°C for 24 h, as described by Murray (1992).

### Cell envelopes isolation and chromatography

Cell envelopes were isolated at 4°C in dim light as described in Farci *et al.*, (2021). Briefly, after harvesting by centrifugation (5000 g, 10 min, 4°C), cells were resuspended in a 50 mM sodium phosphate buffer at pH 7.8 (buffer A) supplemented with DNase I (100 U, DNase I recombinant, RNase-free Roche), and disrupted using a French Pressure Cell (3 cycles at 1100 psi). The sample was centrifuged twice (2000 g, 10 min, 4°C) to remove the debris, then the supernatant was centrifuged to collect cell envelope fragments (48000 g, 10 min, 4°C). The final pellet was resuspended in buffer A, digested with 100 μg/mL lysozyme (8 h, 30°C, under shaking at 800 rpm) to remove surface polysaccharides. Digested cell envelope fragments were centrifuged (48000 g, 10 min, 4°C), and washed three times with buffer A by a serial sequence of centrifugation/resuspension steps. Final samples were used for the cryo-electron crystallography and tomography.

Anionic exchange chromatography was performed according to Farci *et al.*, (2021). Briefly, cell envelope fragments were resuspended (3-5 mg/mL total proteins) and solubilised for 30’ at room temperature with a final concentration of 1.1% of n-dodecyl-β-D-maltoside (β-DDM). After solubilization and centrifugation (48000 g, 10 min, 4°C), the supernatant was subjected to anion-exchange chromatography (Hi-load HP column, Amersham). After 5 column volumes of washing with buffer B (50 mM sodium phosphate, pH 7.4; 0.05% (w/v) β-DDM) at a flow rate of 0.5 mL/min, the elution was done with a linear gradient of 0-2.5M NaCl in buffer C (50 mM sodium phosphate, pH 7.4; 2.5 M NaCl; 0.05% (w/v) β-DDM). All chromatography columns were subjected to the ReGenFix procedure (https://www.regenfix.eu/) for regeneration and calibration prior to use.

### Electron Microscopy

#### Cryo-electron microscopy

The grid preparation and data acquisition were done at CEITEC, Brno, Czech Republic. Quantifoil R2/1.3 holey carbon grids were glow-discharged prior to use. Cell envelope samples were vitrified using a Vitrobot plunge-freezing machine (Mark IV, ThermoFisher) at room temperature (blot force 2, blotting time 2 s, 100% humidity), and placed in autogrid (FEI, Eindhoven, Netherlands) prior to image acquisition. Grids were transferred to a Titan-Krios TEM (ThermoFisher) operating at 300 kV and equipped with a Cs-corrector (cs 2.7 mm), a Quantum GIF energy filter (slit width set to 20eV), and a post-GIF K2 camera (Gatan) at a magnification corresponding to a pixel size of 2.28 Å/px (~ 21000x). A dose-symmetric scheme (3° increment, range ± 60°) was used for tomography acquisition by SerialEM software (Mastronarde, 2005). The defocus was set to vary between 2 μm and 5 μm.

The isolated protein complexes were negatively stained on glow-discharged copper grids (EMS200-Cu) covered with a 20-nm carbon film. The samples were applied at the grid in a volume of 3 μL and the excess was removed after 1 min by capillarity through a delicate touching of the grid bar with a filter paper (Whatman No. 1) for 2 sec. The grids were then stained for 1 min with 5 μL of 2% UranyLess TEM staining solution (Micro to Nano) and the excess of staining was removed by capillarity as in the previous step. Micrographs were acquired with a Tecnai F20 microscope (ThermoFisher Scientific) operating at 200 kV, with an FEI Eagle 4K CCD camera, at a magnification of 53000 x.

#### Cryo-Electron Crystallography processing of cell envelopes fragments

We collected 42 tomograms in movie mode (each tilting angle has a movie of 8 frames) for which CTF and ice quality were inspected. A total of 40 tomograms were selected for further processing. CTFs and resolution were estimated by CTFFIND3 (Mindell and Grigorieff, 2003). For cryo-electron crystallography, 8-frames movies were drift corrected using MotionCor2 (Zheng *et al.*, 2017) on the Focus package (Biyani *et al.*, 2017). Micrographs, at different tilting angles (see Table 1 for tilt angle distribution) and from the 6 best-diffracting tomograms, were processed on the same package but this time in 2D crystallography mode according to the latest implementations of the *2dx* package (Gipson *et al.*, 2007). By this procedure, it was obtained a complete set of information for each micrograph (defocus, tilt geometry, lattice, phase origin) and used to build a 2D projection map. Finally, data from the 73 best images were merged to obtain the final projection map by using the merge suite from Focus in 3D crystallography mode. The 3D model’s visualization and the fittings were done using the Chimera software (Pettersen *et al.*, 2004).

#### Cryo-electron tomography and subtomogram average processing of cell envelopes fragments

Tomograms were processed by the *etomo* software (Mastronarde, 2005). Gold particles were used as fiducial markers and only tomograms with minimal alignment errors were selected for the final reconstruction (Mastronarde, 2005). For sub/tomogram averaging on 3D reconstructions, sub-tomograms boxed with a size of 300×300×300 voxels were manually selected and extracted. Sub-tomogram alignment and averaging with missing-wedge compensation was performed with the PEET software (Cope *et al.*, 2011; Heumann *et al.*, 2011) as follows. Sub-tomograms were initially rotationally aligned assuming that translational shifts of sub-tomograms were approximately correct. Eulerian angles were determined by systematic search over some specified range of values with the range and the coarseness of the search being reduced in successive iterations of the search (Bostina *et al.* 2007). Absolute values of cross-correlation were used for alignment potentially helping to prevent noise by reinforcing to match features in the reference (Cope *et al.*, 2011; Heumann *et al.*, 2011). Finally, p6 symmetrization was performed using the scripts available for PEET. Isosurface visualization was accomplished using the Chimera software (Pettersen *et al.*, 2004).

#### Data and code availability

The final 3D volumes and other relevant information about data acquisition and processing have been deposited in the Electron Microscopy Data Bank, EMD-14095, EMDB-14096; EMDB-14097. All data will be released upon publication. This paper does not report any original code. Any additional information required to reanalyze the data reported in this paper is available from the lead contact upon request.

### Quantification and statistical analysis

Cell envelopes isolation for data analyses was performed on more than 20 independent preparations. Micrographs, tomograms, and maps are representative of at least three independent replicates. Electron crystallographic image statistics and main images parameters are indicated in Table 1. All attempts to reproduce the results here presented were successful.

### Reagents and Tools Table

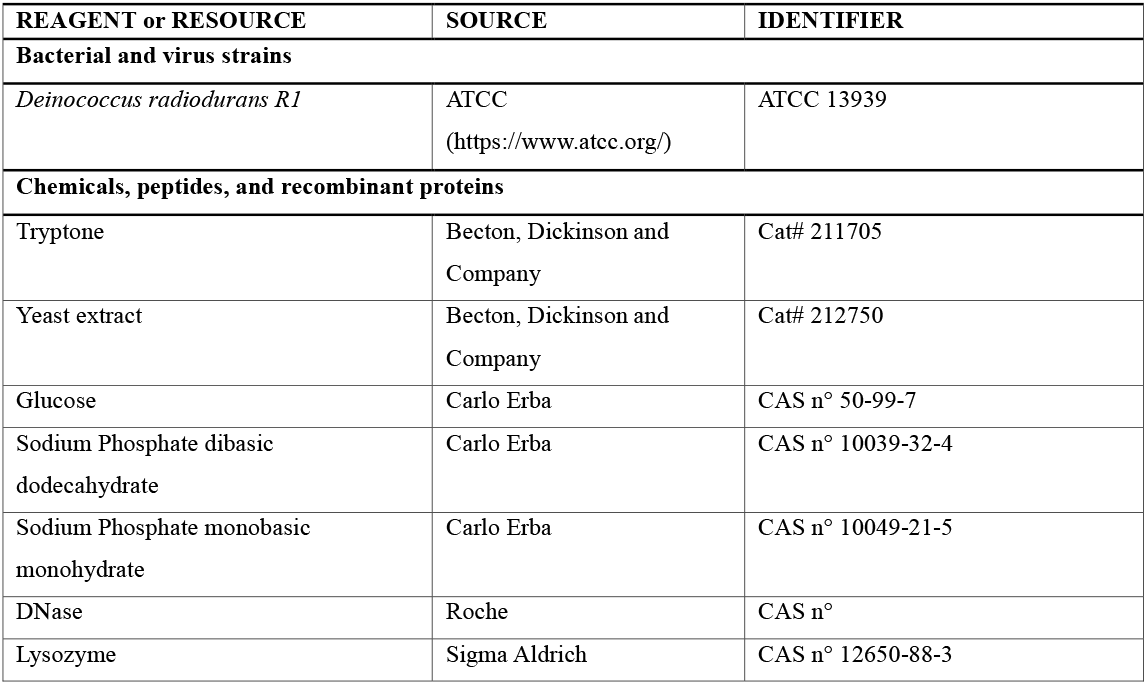
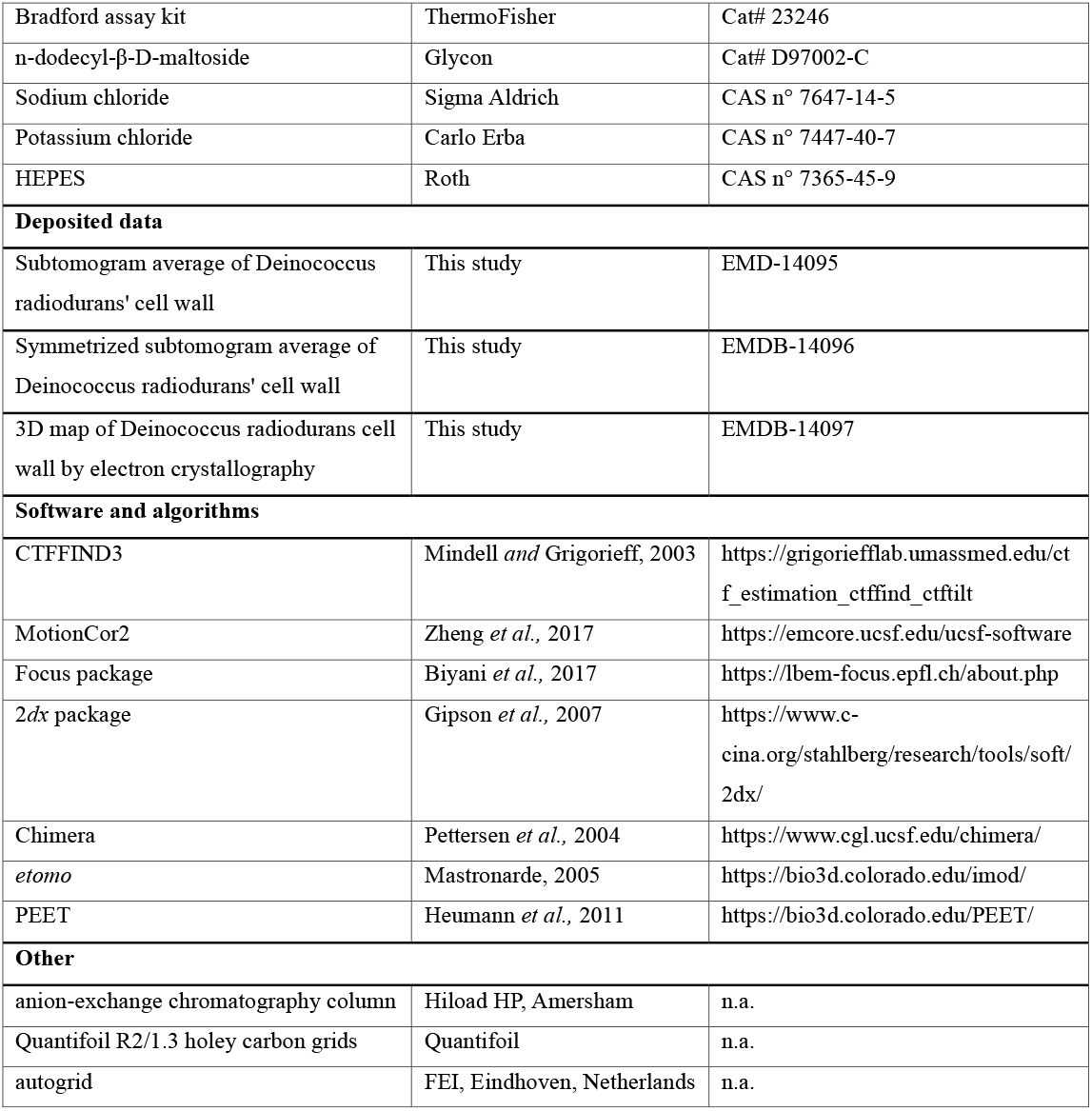

## Acknowledgments

This work was supported by the National Science Center (Poland) with the Sonata BIS 7 Program (2017) Grant PRO-2017/26/E/NZ1/00344 and the Harmonia 10 Program (2018) Grant PRO-2018/30/M/NZ1/00284 (both to D.P., D.F., and P.H.). We acknowledge the Cryo-electron microscopy and tomography core facility CEITEC MU of CIISB, Instruct-CZ Centre supported by MEYS CR (LM2018127). DF kindly thanks Dr. J.M. Heumann (University of Colorado Boulder) for the support with the subtomogram averaging processing using PEET.

## Author contribution

Conceptualization: DP, DF. Investigation: DP, DF, PH. Visualization: DP, DF. Funding acquisition: DP. Project administration: DP, DF, PH. Supervision: DP. Writing – original draft: DP, DF, PH.

## Inclusion and Diversity

While citing references scientifically relevant for this work, we actively worked to promote gender balance in our reference list. The author list of this paper includes contributors from the location where the research was conducted who participated in the data collection, design, analysis, and/or interpretation of the work.

## Supplemental figures and videos

**Supplementary Figure 1: Tomographic parameters and resolution data.** In the image is reported the Fourier Shell Correlation (FSC) for the symmetrized subtomogram average; the dashed-red line represents the spatial frequency with cutoff at 0.5 (Nyquist frequency - gold standard). The inset table reports the CTF parameters. Related to Figure 6.

**Supplementary Figure 2: Side view of the symmetrized map obtained by subtomogram average.** Map showing the T4P-like complex occupancy (in orange) and its position with respect to the S-layer (SL), outer membrane (OM), and inner membrane (IM) as observed in the present study (see also Sup. Fig. 3). The extracellular, periplasmic, and cytoplasmic spaces are also indicated. The scale bar indicates 50 Å. Related to Figure 6.

**Supplementary Figure 3: Cell envelope layering and membranes localization.** On the left is shown the tomographic reconstruction of a patch. The S-layer/outer membrane system (SL/OM) and the inner membranes (IM, black arrows) are highlighted with black arrows. The scale bar indicates 200 Å. On the right, it is shown a detail of the same cell envelope processed by subtomogram average with indicated cytosol, periplasm, and extracellular space. The scale bar indicates 50 Å. The inset table on the bottom summarizes the thickness values of the cell envelope and its regions. The values in the inset table represent the mean of 40 independent tomograms. Related to Figure 5.

**Supplementary Movie 1**: Electron density map showing the features of the diffracting cell envelope fraction with a detail of its main complexes. The T4P-like (orange), the SDBC (pink), and the dihedral complex (yellow) were extracted and refitted into the map. Related to Figure 3.

**Supplementary Movie 2:** Representative tomogram of a cell envelope patch reconstructed from a typical specimen. Related to Figure 5.

