## Supplementary material for "Mesoscale organization in the cell envelope of *Deinococcus radiodurans*": Sup.

### Sup. Figure 1

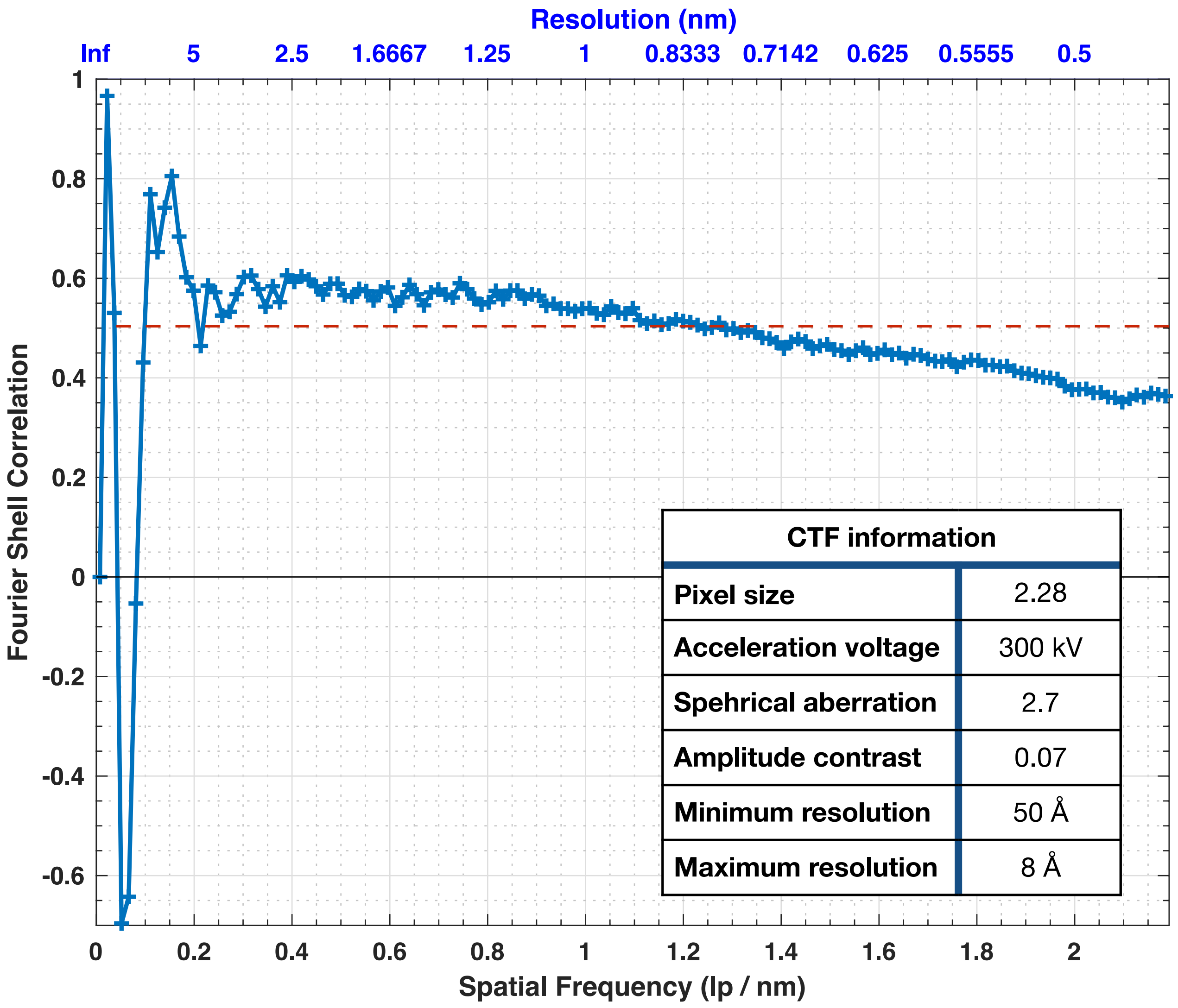


| CTF information | |
|---|---|
| **Pixel size** | 2.28 |
| **Acceleration voltage** | 300 kV |
| **Spehrical aberration** | 2.7 |
| **Amplitude contrast** | 0.07 |
| **Minimum resolution** | 50 Å |
| **Maximum resolution** | 8 Å |

### Sup. Figure 2

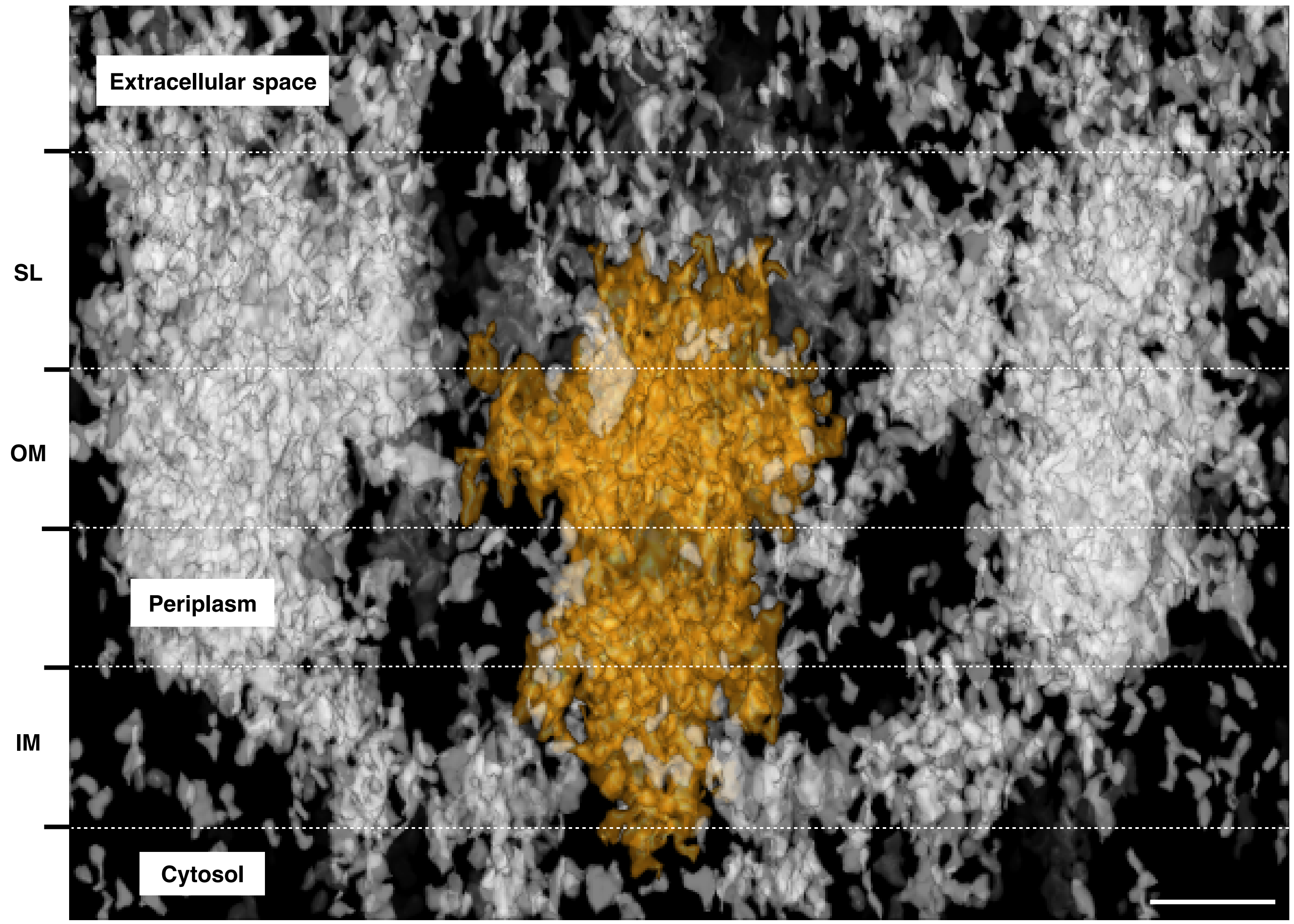

### Sup. Figure 3

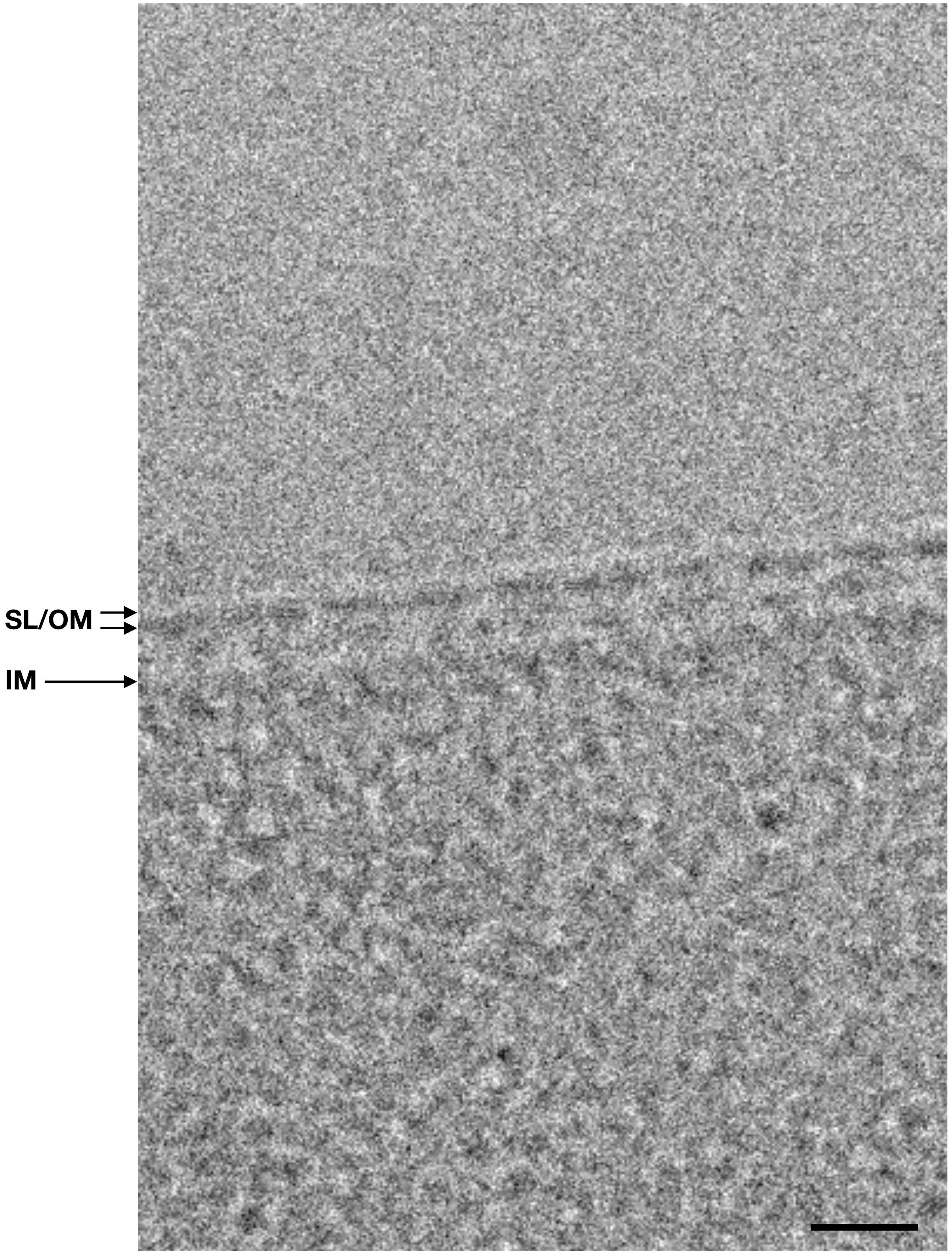


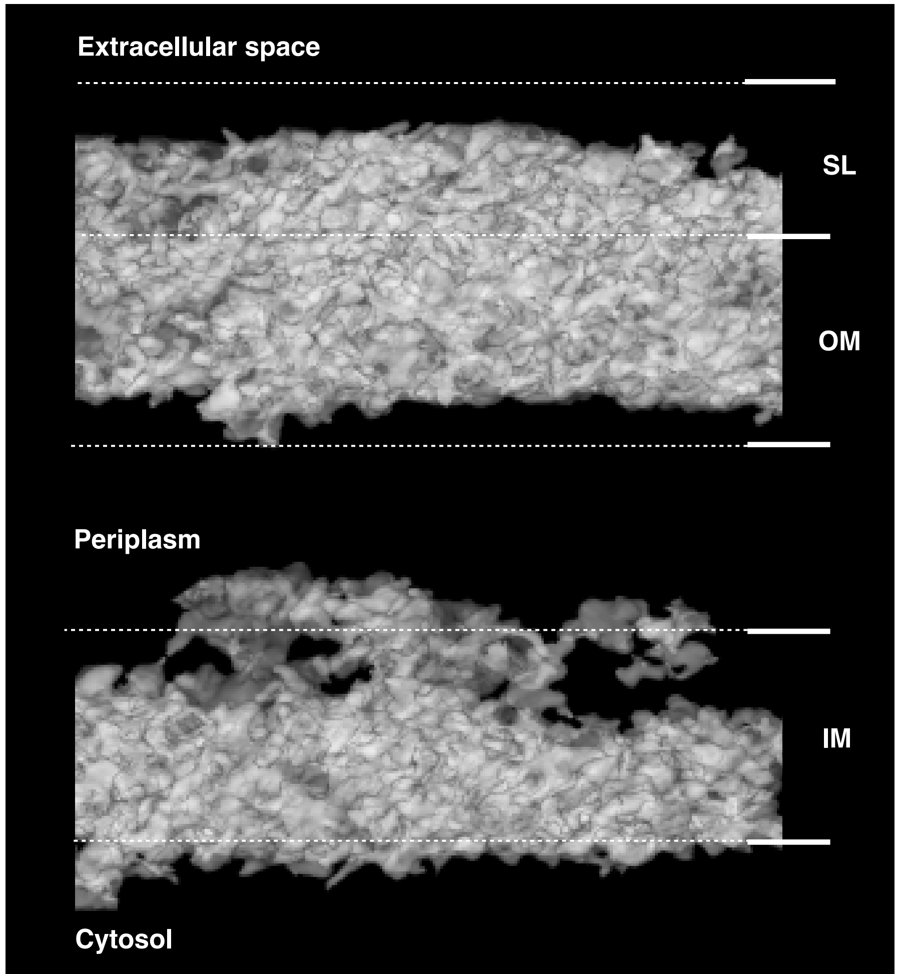


| Membranes thickness | Average (Å) | S.D. (Å) |
|---|---|---|
| **Full membranes** | 299.53 | ± 2.88 |
| **S-layer/Outer membrane** | 98.9 | ± 9.40 |
| **Periplasm** | 99.3 | ± 12.2 |
| **Inner membrane** | 69.8 | ± 5.4 |
